# Shared environments complicate the use of strain-resolved metagenomics to infer microbiome transmission

**DOI:** 10.1101/2024.07.18.604082

**Authors:** Reena Debray, Carly C. Dickson, Shasta E. Webb, Elizabeth A. Archie, Jenny Tung

## Abstract

In humans and other social animals, social partners have more similar microbiomes than expected by chance, suggesting that social contact transfers microorganisms. Yet, social microbiome transmission can be difficult to identify based on compositional data alone. To overcome this challenge, recent studies have used information about microbial strain sharing (i.e., the shared presence of highly similar microbial sequences) to infer transmission. However, the degree to which strain sharing is influenced by shared traits and environments among social partners, rather than transmission *per se*, is not well understood. Here, we first use a fecal microbiota transplant dataset to show that strain sharing can recapitulate true transmission networks under ideal settings when donor-recipient pairs are unambiguous and recipients are sampled shortly after transmission. In contrast, in gut metagenomes from a wild baboon population, we find that demographic and environmental factors can override signals of strain sharing among social partners. We conclude that strain-level analyses provide useful information about microbiome similarity, but other facets of study design, especially longitudinal sampling and careful consideration of host characteristics, are essential for inferring the underlying mechanisms.

## INTRODUCTION

Animals are born sterile and presumably acquire their microbiomes from a combination of vertical, environmental, and horizontal transmission – including, in social species, through social interactions with their conspecifics. In the last ten years, studies in a wide variety of social animals–from humans to birds, insects, and other primates–have shown that social group residency and social network structure have significant explanatory power for the composition and genetic content of animal microbiomes, especially in the gut^1–5^. These observations suggest that social transmission may be an important mechanism shaping animal microbiomes, with potential consequences for microbiome stability and diversity, host infectious disease risk and immune function^6,7^, and the evolution of host behaviors to facilitate and/or avoid microbial transmission. From the perspective of microorganisms, social transmission may also generate selection pressures to specialize on the substrates and conditions available in the host environment, causing the fitness of socially transmitted microbes to become more closely aligned with the fitness of their hosts^8^. Indeed, an early hypothesis proposed that some social behaviors, such as trophallaxis and coprophagy, evolved in part to transmit specialized, beneficial microbes that can no longer tolerate environmental intermediates^9,10^.Consistent with this idea, several studies in the more recent era of microbiome sequencing report that anaerobic and non-spore-forming species (i.e., those with low predicted environmental persistence) are more likely to be socially shared^2,11,12^.

Together, these studies argue that social transmission, either directly between interacting animals or indirectly through intermediate substrates (e.g., shared nesting material, deposited scent marks), may play an important role in host microbiome structure and function. This idea is bolstered by experimental work: for example, evidence from captive populations of rhesus macaques, goats, pigs, and vampire bats all show that moving individuals from separate to shared housing causes their microbiomes to converge^13–15^. However, although social transmission has become a favored explanation for socially structured microbiomes, it can be very difficult to distinguish from alternative mechanisms also at play in natural populations. Many microbial species are widely distributed in nature^16^. Individuals may therefore independently acquire lineages of the same species, especially if they share similar environments. For example, many animals forage in groups with, or share food preferentially with, close social partners, leading to correlations between social group membership, the strength of social bonds, and diet^17,18^. Social groups may also encounter microbial species in the water, soil, and other substrates in the environment that differ based on territory or space use^19^. In some species, animals associate preferentially with their close genetic relatives and/or with individuals of a similar age^20,21^. If these characteristics affect which microbial species can establish and persist in hosts, then age, kin, or shared environmental effects could be mistaken for social transmission, inflating estimates of social effects on the microbiome^22,23^. Few (if any) studies in natural populations, including humans, can completely eliminate these alternative explanations, and direct experimental evidence for transmission is difficult to obtain. Consequently, the relative importance of social transmission in explaining socially structured microbiomes remains an open question.

One proposed solution is to leverage computational approaches that classify microbial reads at subspecies or strain levels from shotgun metagenomic data^24–26^. Most current strain profiling pipelines either use sequence variation in species-specific marker genes to construct strain-level phylogenies^24,25,27^, or align short reads to a set of reference microbial genomes to identify variants throughout the genome^26,28^. Samples can be classified as ‘sharing a strain’ if the lineages they carry are close in the phylogeny^29^, have highly similar marker gene sequences^25^, or have high genome-wide nucleotide identity^26,30^. Individuals with a higher proportion of strains in common are then typically assumed to engage more frequently in transmission. Strain profiling in humans has revealed elevated strain sharing rates among mothers and infants^31–33^, interpreted as vertical transmission, and among household members^28,34^ and village residents^29,35^, which have been interpreted as support for pervasive horizontal transmission through social networks. Strain-resolved approaches have not yet been widely applied in natural populations of other social animals, but early evidence in wild baboons also demonstrates elevated rates of strain sharing within versus between social groups^36^, suggesting that findings in humans will likely generalize to other host species.

Elevated strain sharing rates are often assumed to be the direct result of transmission, based on the assumption that individuals are otherwise unlikely to acquire the same strain independently^34,37^. Consequently, socially structured strain sharing has been treated as evidence for the importance of direct person-to-person transmission. But the validity of this assumption is unclear. Some pairs of individuals might frequently transmit strains among each other, but consume different enough diets (for instance) that they retain little of what they receive. Others may exchange strains only occasionally but retain a high proportion of socially acquired strains, resulting in similar microbiomes overall. The dual processes of transmission and retention/persistence make it difficult to evaluate whether strain sharing rates are elevated among social partners because they truly exchange microbes, or simply because they experience similar environments that shape their microbiomes in parallel.

Here, we evaluate the explanatory power of strain sharing rates in two gut microbiome data sets: an experimental fecal microbiota transplant with a known underlying transmission network, generated by Ianiro and colleagues^38^, and a wild baboon population where social structuring of the gut microbiome has been described in previous work^2^. The baboon population has been the subject of continuous study by the Amboseli Baboon Research Project for over 50 years, and extensive data are available on the behavior, ecology, life histories, and gut microbiomes of individually recognized study subjects, making it an ideal setting for disentangling drivers of microbial strain sharing^39^.

We use the fecal microbiota transplant to first ask whether, in a setting in which the transmission network is completely understood, strain-level resolution improves our ability to infer the transmission network compared to coarser, compositional resolution. We test whether species that follow the true transmission network are more likely to come from specialized, host-associated taxa, as predicted by evolutionary theory^9^. We also take advantage of this data set to evaluate alternative criteria for defining a strain sharing event as a case of true transmission. We find that transmission dynamics can be more reliably resolved in strains that are private to a single individual at an earlier time point and then spread to other individuals, whereas strains that are already widespread provide less robust information about transmission^11^. Using the baboon data set, we then consider cases of strain sharing under natural conditions – the setting of greatest interest in studies of social transmission^40^. We evaluate “background” rates of strain sharing among individuals with non-overlapping lifespans (i.e., cases where social transmission is impossible) and compare those rates to both strain sharing levels in longitudinal samples from the same individual and close social partners. Finally, we test whether environmental or demographic characteristics provide alternative explanations to social transmission. Together, our analyses suggest that, although strain-resolved metagenomics has substantial value for understanding microbial transmission in the gut microbiome, elevated strain sharing rates are, by themselves, insufficient to infer direct social transmission.

## RESULTS

### Strain sharing in a known transmission network

To assess whether bacterial strain sharing rates can act as a reliable indicator of microbiome transmission, we first identified a sample set with known transmission dynamics. Here, we drew on fecal metagenomic sequences from a fecal microbiota transplant (FMT) study, which tracked both healthy donors and FMT recipients (patients with inflammatory bowel disease or chronic *Clostridioides difficile* infection) living in Rome, Italy^38^. Each patient underwent a three-day vancomycin regimen and then received a transplant prepared from the fecal sample of a single donor. Stool samples were collected from donors immediately before FMT, and from recipients immediately before FMT and 15-30 days after FMT (n=21 total samples, from 13 subjects, including 5 healthy donors and 8 FMT recipients, Figure **1a**). This study design allowed us to compare strain sharing in actual donor-recipient pairs (“matched pairs”) to background rates of strain sharing in all other pairs of “mismatched” donors and recipients (Figure **1b**).

**Figure 1.**
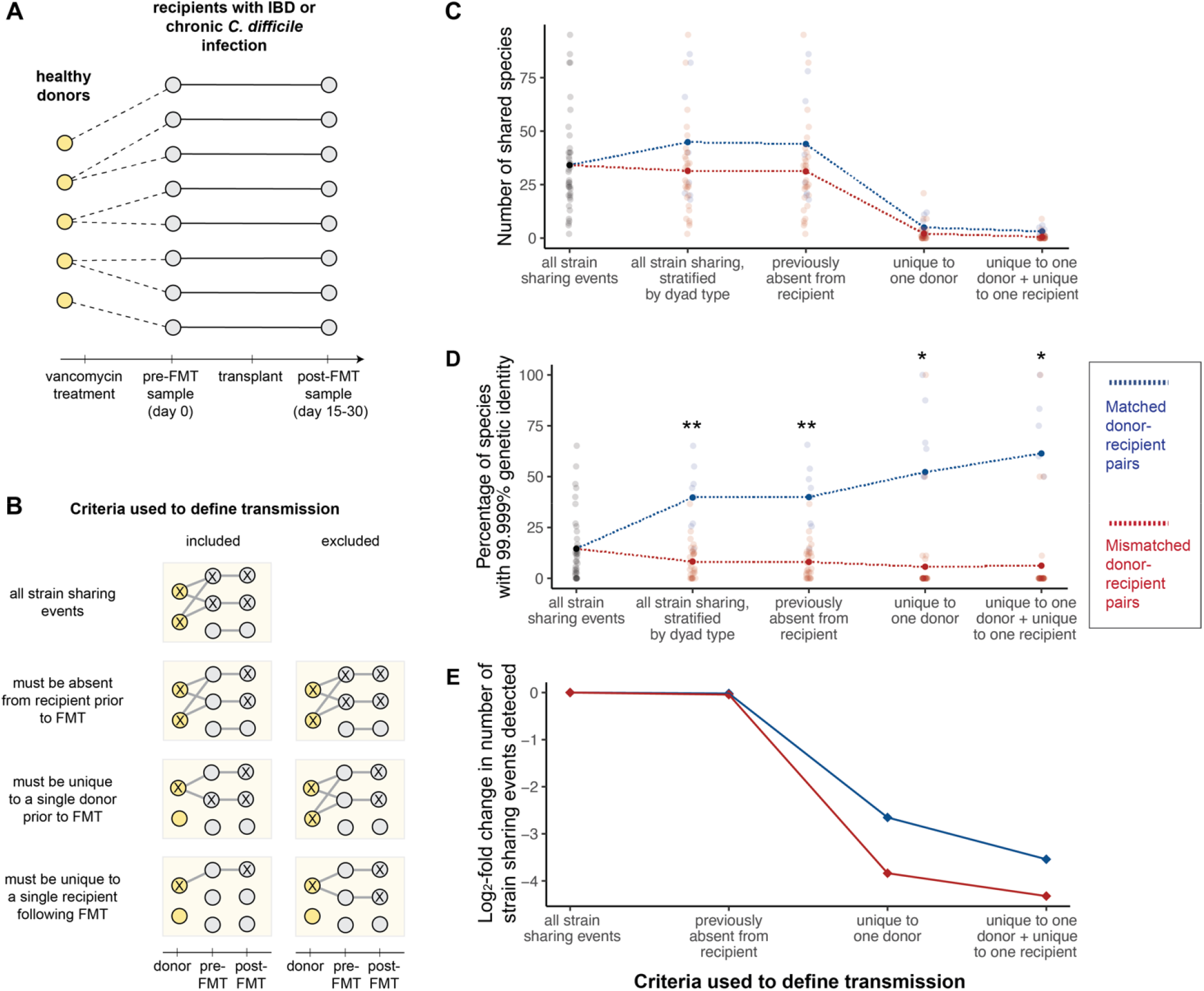
Strain sharing among donors and recipients in a fecal microbiota transplant cohort. **(a) Transmission network in the FMT study.** Stool samples from five healthy donors and eight recipients enrolled in a clinical study were collected for metagenomic sequencing and analysis^38^. **(b) Visualization of criteria used to define a putative transmission event**. Each box shows examples of strain sharing events that would be considered transmission (included) or not considered transmission (excluded) based on increasingly stringent criteria concerning the prevalence of the strain in the population. All criteria were cumulative; e.g., the list of strain sharing events absent in pre-FMT recipients was used as the starting point to further select strain sharing events that were unique to a single donor. Inferred transmission events under each successive set of criteria are shown using solid lines to connect samples; x’s mark the presence of shared strains. **(c) Species sharing across increasingly stringent definitions of transmission**. The number of shared microbial species among matched donor-recipient dyads (blue) and all other donor-recipient comparisons (mismatched: red) is shown following serially more stringent filtering criteria. **(d) Strain sharing across varying definitions of transmission**. The percentage of strain sharing events among matched donor-recipient dyads (blue) and all other comparisons (red) is shown following serially more stringent filtering criteria. Asterisks represent significant differences between matched and mismatched cohorts based on a t-test followed by Benjamini-Hochberg correction: (***) p<0.001; (**) 0.001≤p<0.01; (*) 0.01≤p<0.05. **(e) Subsets of strain sharing events detected under varying criteria for transmission**. The log-scaled proportion of total strain sharing events that were classified as transmission under varying criteria. Strain sharing between matched pairs is represented in blue and strain sharing between mismatched pairs is represented in red.

We measured strain sharing using the inStrain pipeline, which calculates the average nucleotide identity (ANI) of bacterial species shared between two individuals^26^. We considered pairs of donors and recipients to share the same strain of a species if their respective sequence alignments met or exceeded a threshold of 99.999% ANI. This threshold was set based on developer recommendations and is estimated to discriminate between strains that diverged within the last 2.2 years^26^. We first calculated strain sharing rates for all microbial species and in all post-FMT donor-recipient pairs (both matched true donor-recipient pairs and mismatched pairs). We identified cases of strain sharing for 131 of the 755 species we detected in the data set. An average of 15% of species shared between a given donor-recipient pair (matched or mismatched) met the sequence identity threshold to be considered the same strain. Consistent with our expectations, strain sharing rates were significantly higher among matched donor-recipient pairs (i.e., true links in the underlying transmission network) than mismatched pairs, even though matched pairs did not share significantly more *species* in common (strain sharing: 40% between matched donor recipient pairs versus 8% between mismatched pairs, Benjamini-Hochberg adjusted p=0.002; species sharing: mean=44.9 shared species in matched pairs versus 31.4 in mismatched pairs, adjusted p=0.253; Figure **1c,d**).

We then asked whether imposing more stringent criteria on strain sharing events would improve our ability to differentiate matched from mismatched pairs, and therefore the concordance between the strain sharing network and the true underlying transmission network. First, for each pair (whether matched or mismatched), we excluded strains shared prior to the FMT from further consideration. This approach controls for background rates of strain sharing between donors and recipients, but had little effect on strain sharing patterns in practice, suggesting that background strain sharing between donors and recipients was generally low (Figure **1d**, “strains previously absent from recipient”). Next, we reasoned that sharing of strains that were rare in the data set might be more likely to reflect true transmission, whereas sharing of widespread strains is more likely to represent independent acquisition. Further restricting the dataset to exclude strains detected in more than one donor *prior* to FMT was more effective at distinguishing the true transmission network. Although the number of shared species under consideration decreased with this filter (Figure **1c**), the *proportion* of shared species with 99.999% or higher ANI increased to 52% in matched donor-recipient pairs, compared to only 6% in mismatched pairs (adjusted p=0.01; Figure **1d**, “strains unique to one donor”). Further restriction of the dataset to additionally exclude strains that were detected in more than one recipient *after* FMT moderately improved this result (61% between matched pairs versus 7% between mismatched pairs, adjusted p=0.01, Figure **1d**, “strains unique to one donor + unique to one recipient”).

Naturally, the increasingly stringent criteria for transmission excluded strain sharing events among matched donor-recipient pairs as well. For example, the most stringent threshold excluded ∼95% of strain sharing events in mismatched donor and recipient pairs from being categorized as true transmission, but also excluded ∼91% of strain sharing events that occurred in correct pairs (Figure **1e**). The focus on strains that are unique in donor and/or recipient cohorts is therefore useful for obtaining the clearest representation of the true transmission network, but more relaxed criteria may be appropriate for studying other aspects of strain sharing in a broader and more representative set of gut microbial species.

### Candidate transmission events in FMT are enriched for rare, anaerobic, and host-specific bacterial taxa

Having established that matched donors and recipients shared *more* bacterial strains than mismatched pairs, we next asked whether they shared *different* bacterial taxa as well. Species sharing events in matched donor-recipient pairs involved subtly different sets of bacteria taxa than species sharing events in mismatched pairs (Fisher’s exact test, p=0.024; Figure **S1a**). Species from the Erysipelotrichaceae family were enriched in matched pairs compared to mismatched pairs (log_2_ odds ratio=2.73, p<0.001, Figure **S1b**). Strain sharing events among matched and mismatched pairs also involved different sets of bacteria taxa, though no families were sufficiently enriched among matched pairs to detect in family-by-family analyses (Fisher’s exact test, p=0.048; Figure **2a**, Figure **S1c**).

There was a slight difference in the initial population prevalence of strains shared in matched versus mismatched pairs, with strain sharing in matched pairs coming from rarer species (t=2.36, p=0.019; Figure **2b**). Strains shared by matched pairs were also more likely to come from obligately anaerobic genera (permutation test, p<0.001, Figure **2c**), and the proportion of strain sharing events involving obligately anaerobic taxa increased across progressively stricter definitions of transmission. Similarly, with stricter criteria for transmission, strain sharing events were increasingly likely to involve bacterial genera that were reported in human-derived microbiome samples, but not in any other host species, in the Genomes OnLine database^41^. Under the strictest transmission criteria, for example, 78% of strain sharing events involved human-specific taxa, compared to only 31% in the strain sharing dataset as a whole (permutation test; unique to one donor, p=0.002; unique to one donor and recipient, p=0.007; Figure **2d**).

**Figure 2.**
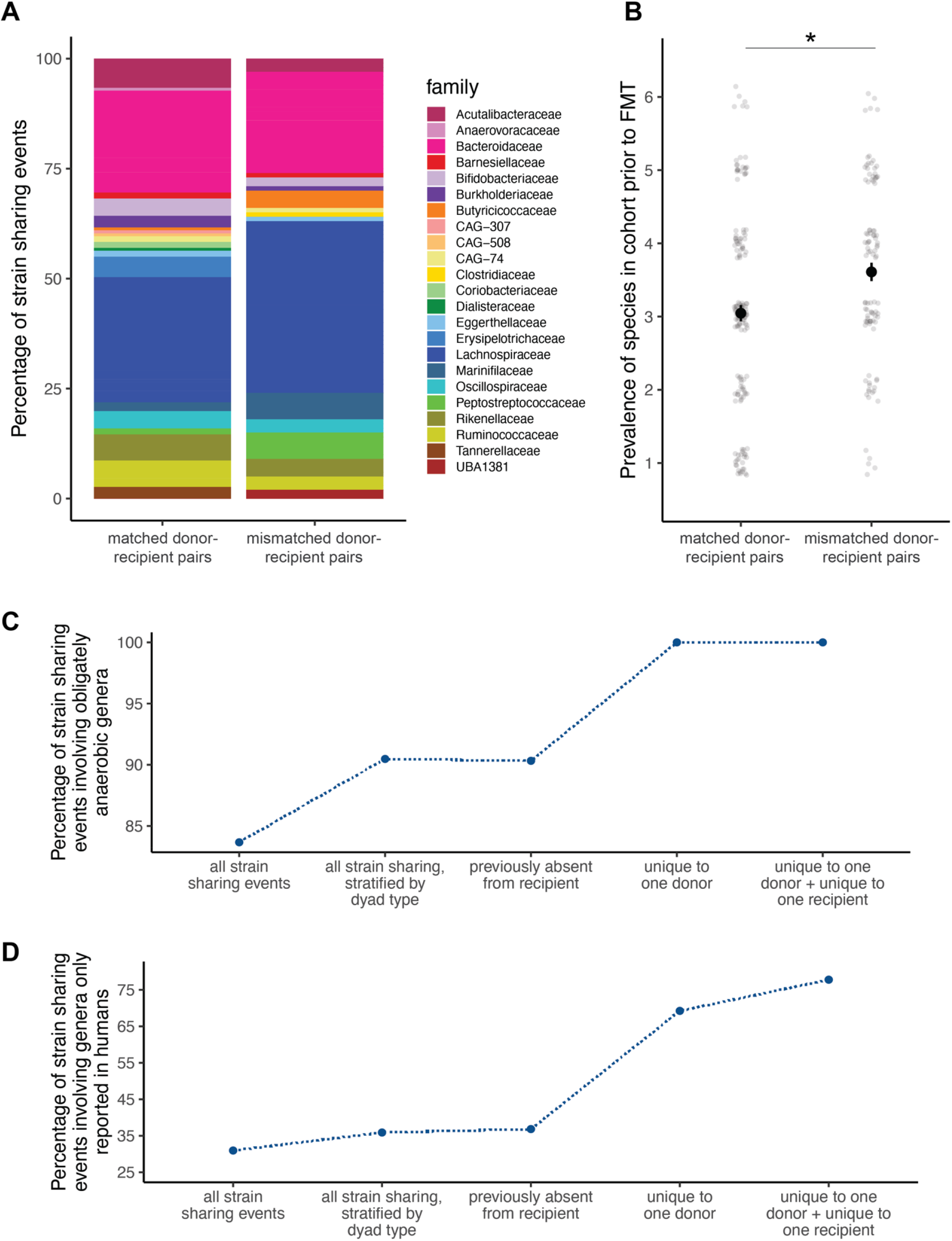
Taxonomic and functional characteristics of bacteria that exhibit strain sharing between donors and recipients of fecal microbiota transplants. **(a) Family-level structure of bacterial taxa that were shared at the strain level between subjects**. Bar chart of species, grouped into families, that exhibit strain-sharing between matched (left) or mismatched (right) pairs, based on a 99.999% ANI threshold. Colors reflect different bacterial families and are scaled to represent the relative proportions of strain-sharing events in each category. **(b) Population prevalence in donors and pre-FMT recipients**. Each strain sharing event was annotated according to the initial prevalence of the species to which it belonged. The y-axis represents the number of donors or pre-FMT recipients containing the species (of a maximum possible of 13); points are jittered along both x and y axes for better visibility. **(c) Anaerobic metabolism across varying definitions of transmission**. Each strain sharing event between matched pairs was annotated, at a species level, as aerobic, anaerobic, or mixed. **(d) Host specificity across varying definitions of transmission**. Each strain sharing event between matched pairs was annotated as a genus reported only in humans or reported in multiple species.

### Strain sharing in a natural primate population

The FMT data show that strain sharing patterns can, in principle, reflect the underlying transmission network in a short-term, experimental cohort. However, the primary interest in strain sharing across social and/or transmission networks focuses on unmanipulated groups or populations, where strain sharing may arise due to transmission, genetic effects, and/or shared environments. We therefore next asked whether strain sharing is a reliable indicator of transmission in natural populations, using fecal samples collected from baboons in the Amboseli ecosystem of Kenya^42^.

In the Amboseli baboon population, as in any natural setting, there is no “known” transmission network. However, we reasoned that strain sharing due to transmission should be more likely among some pairs of individuals than others. For example, baboons who live in different social groups throughout their entire lives would have fewer opportunities for microbial transmission than close social partners who interact frequently. To test this prediction, we generated metagenomes from a set of 126 fecal samples collected from 93 individual baboons between 2007-2017 (mean sequencing depth=37 million read pairs ± 22 million s.d.; Figure **3a**). We compared: i) 23 pairs of baboons whose lives never overlapped; i.e., the first one died before the second was born; ii) 20 pairs of baboons whose lives overlapped, but lived in different social groups the entire time; and iii) 26 pairs of baboons that were close social partners living in the same social group and sampled within 4 days of each other. Because high rates of strain sharing are expected from samples collected from the same individual over time^43,44^, relative to the first three categories, we also included iv) 22 pairs of longitudinal samples from the same individual, collected 4-5 months apart.

**Figure 3.**
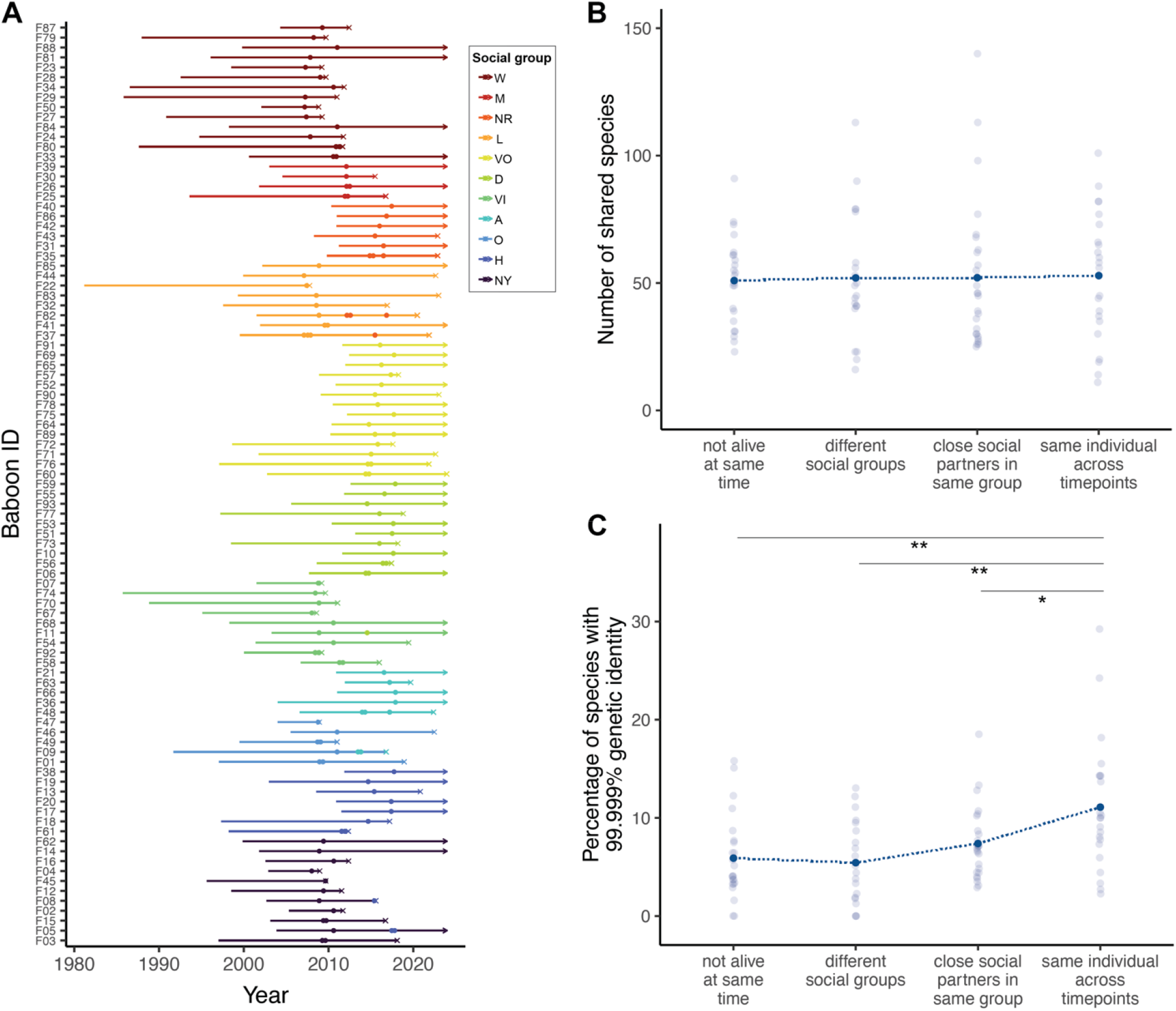
Species and strain sharing rates in the gut microbiomes of wild baboons. **(a) Individuals included in this dataset.** Each line represents an individual baboon, starting from the year it was born. The line terminates either in an (x) to represent death or an arrow to represent that the animal was alive as of December 31st, 2023. Points represent sampling events. Segments are colored according to social group membership when the individual was born, while points are colored according to social group membership when the corresponding samples were taken. As females do not typically disperse in this species, animals that belonged to two or more groups during their lifetimes represent group fission or fusion events. **(b) Species sharing across dyad types**. The number of shared species between each dyad based on inStrain profiling (95% popANI). **(c) Strain sharing rates across dyad types**. The percentage of shared strains between each dyad (99.999% popANI). Only repeated samples from the same individual differed from any of the other categories (Tukey HSD; same individual - close social partners p=0.045, same individual - different social groups p=0.001, same individual - not alive at same time p=0.0027).

If co-residency and social interactions facilitate transmission of gut microbes, then species and strain sharing should be low in categories (i) and (ii) and higher in category (iii). In contrast to this expectation, there were no systematic differences in species sharing across categories (Figure **3b**). Within shared species, strain sharing rates also did not differ among baboons whose lives never overlapped (5.9%), baboons who lived in different social groups (5.4%), and baboons who lived in the same social group and interacted closely (7.4%). Only repeated samples from the same individual had significantly elevated strain sharing rates compared to the other categories (11.1%; Figure **3c**). There were also no differences in the family-level taxonomic composition of strains shared across categories (Fisher’s exact test, p=0.463, Figure **S2a**) or the proportion of strain sharing events involving anaerobic bacteria (Fisher’s exact test, p=0.353, Figure **S2b**). Strain sharing events among close social partners were more likely to involve rare species compared to strain sharing across social groups (Tukey HSD, p=0.035), but not when compared to baboons that lived at different times entirely (Tukey HSD, p=0. 145, Figure **S2c**).

These results were somewhat surprising in light of the strong tendency to interpret strain sharing as evidence for direct transmission, or specifically social transmission^34,37^. We therefore asked whether similarity in other characteristics, such as age, diet, or time of sampling, inflated strain sharing among baboons who never interacted directly. Among baboons who lived in different social groups at the same time, those with more similar diets in the year they were sampled had more shared strains (linear regression; slope=19.83, df=18, p=0.004; Figure **4a**). This relationship was directionally similar, though weaker and not statistically significant, when diet data were aggregated by month of sampling rather than year (linear regression; slope=3.95, df=18, p=0.388; Figure **S4a**). Baboons had elevated strain sharing rates if they were sampled at similar times of year as well (linear regression, slope=-1.31, df=18, p=0.017; Figure **4c**), possibly due to seasonal variation in their diets and environments (Figure **S4b**).

Among baboons whose lives never overlapped, pairs shared more strains if they were sampled during rainier months (considering strain sharing across both dominant and minor strains; linear regression; slope=0.065, df=21, p=0.009; Figure **4b**). Notably, this pattern was undetectable when strain sharing was based only on sharing the same dominant strain of each species (linear regression, slope=0.002, df=21 p=0.791; Figure **S3**). During rainy periods, baboons therefore appear to be more likely to harbor multiple strains of the same species and share more non-dominant strains.

**Figure 4.**
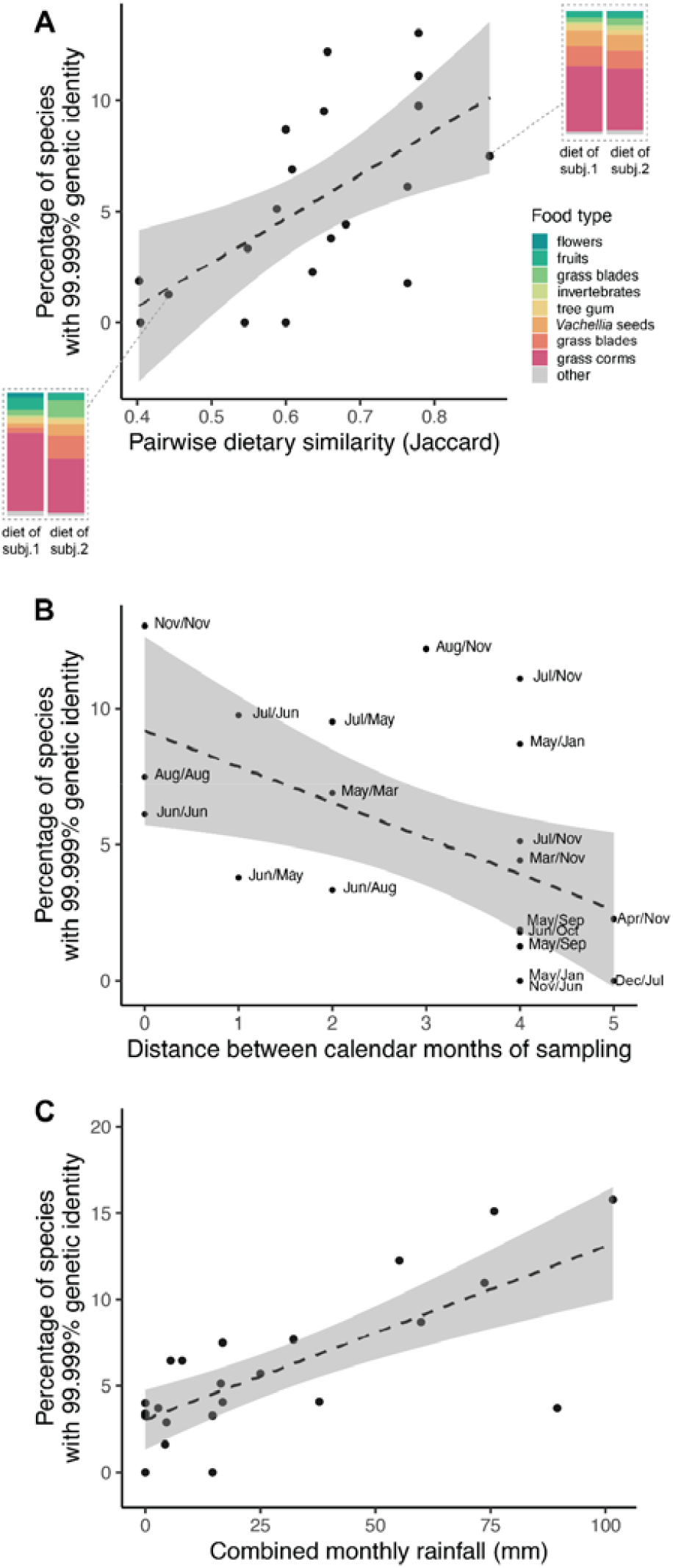
Diet, sample month, and rainfall predict strain sharing among non-coresiding baboons. **(a) Dietary similarity predicts strain sharing.** Dietary similarity was calculated using the Jaccard similarity index based on the dietary compositions of each pair of baboons that lived in different social groups at similar times. Insets display dietary data from selected baboon pairs on either x-axis extreme. **(b) Time of year predicts strain sharing**. Minimum distance between the numeric representations of the months (1=Jan, 2=Feb, etc.), regardless of the year in which they were sampled, for pairs of baboons that lived in different social groups at similar times. No animals from this set were sampled in directly opposing months (i.e., distance=6). **(c) Rainfall predicts strain sharing**. Rainfall was measured daily using a rain gauge and summed by month for samples taken from baboons that lived at different times.

Neither the number of years between the two sampling events nor the age difference between individuals at the times of sampling predicted strain sharing levels (years between sampling: linear regression, slope=-0.424, df=21, p=0.532; age difference: linear regression, slope=0.076, df=67, p=0.424; Figure **S5a**,**b**).

### Concordant findings across two strain sharing pipelines

Our primary approach above uses average nucleotide identity across the observed fraction of each species’ genome to measure strain sharing^26^. An alternative method is to construct a phylogeny based on a set of clade-specific marker genes and classify the closest tips of the phylogeny as the “same strain” rather than using an absolute distance measure^27^. To assess whether these alternative approaches influence strain sharing patterns in our datasets, we therefore also aligned metagenomic reads to the MetaPhlAn marker gene database^45^, then used StrainPhlAn^27^ to build phylogenetic trees for all bacterial species present in four or more individuals. Following previous work, individuals with normalized branch lengths less than or equal to 0.1 (i.e., the 10th percentile of pairwise distances for the species) were considered to share a strain^33^.

In the fecal microbiota transplant dataset, per-dyad estimates of strain sharing were highly correlated between inStrain and StrainPhlAn (Pearson’s correlation, r=0.901, p<0.001 for pairs with 3 or more shared species; Figure **S6a**). Matched donor-recipient pairs shared significantly more strains than mismatched pairs (adjusted p=0.040; Figure **S6b**). The additional filtering criteria were not useful for further resolving the transmission network with StrainPhlAn, as the number of shared strains remaining quickly declined to zero for most pairs (Figure **S6b**).

In the Amboseli baboon population, inStrain and StrainPhlAn also generated correlated estimates of dyad-wise strain sharing rates (Pearson’s correlation, r=0.753, p<0.001 for pairs with 3 or more shared species; Figure **S7a**). As before, strain sharing rates did not significantly differ among baboons in the same social group, baboons in different social groups, and baboons whose lives did not overlap. In this case, however, even longitudinal samples did not differ from any of the other categories (Figure **S7b**). This may be the result of poor representation of baboon gut-associated species in the MetaPhlAn/StrainPhlAn database. StrainPhlAn was only able to identify (and therefore compare) an average of 2.4 species per pair of samples and 98 species total. inStrain profiled an average of 52.0 species per pair and 369 species total, likely because it readily accommodates the addition of microbial genomes assembled directly from this baboon population and other non-human primate species (metagenome-assembled genomes or MAGs)^46^.

## DISCUSSION

Strain-resolved metagenomic analysis has been proposed as a way to reliably track microbial transmission through host populations^11,29,37,47^. Tracking transmission is essential to understanding how contact networks and shared surfaces contribute to microbial dispersal, how transmissibility varies among microbial taxa, and how newly arrived strains establish and interact with the resident microbiome following transmission events^40,47^. We found that strain sharing closely correlated with the true transmission network in a cohort of human patients undergoing a fecal microbiota transplant. Strain-level information significantly improved the correspondence to the transmission network compared to species-level sharing, which was not appreciably higher in matched (true) donor-recipient pairs than among other individuals in the study. Focusing on strains that were limited in prevalence further improved correspondence to the transmission network, suggesting that generally widespread strains may be more likely to be independently acquired by processes other than direct transmission.

Yet in many respects, data from microbiota transplants represent a best-case scenario for detecting microbiome transmission. In the data set we analyzed, the patients had taken antibiotics shortly before the transplant, potentially increasing the probability that the transmitted strains would establish and reach detectable levels in the gut community^38,48,49^. The transplant, which occurred through colonoscopy, bypassed normal transmission routes for strains that might have otherwise been poor colonizers^7^. Recipients were sampled shortly after the transplant, possibly allowing detection of short-term or unstable colonizers before they went extinct^50^. Further, the recipients all experienced gut dysbiosis prior to the FMT, and may have therefore been particularly vulnerable to colonization by new bacterial strains^51^. While the FMT data set therefore provides important proof of principle that strain-resolved metagenomics *can* recover true transmission events, it does not show that this method *does* reliably reflect transmission networks in natural populations, where individuals have the simultaneous potential to be donors and recipients at all times.

Our analysis in the Amboseli baboons points to the much higher complexity of this problem. In contrast to the FMT dataset, background strain sharing in this population was often high among individuals that had never co-resided, especially if they shared other environmental characteristics. For example, baboons that spent their lives in different social groups but ate similar diets had strain sharing rates that often exceeded those of close grooming partners living in the same group and sampled within a few days of each other. This result is qualitatively consistent with a large body of work showing that diet predicts gut microbiome species composition^52–54^, but suggests that diet shapes microbiome similarity at an even finer genetic resolution than previously appreciated. Among baboons whose lives never overlapped at all, strain sharing patterns were explained in part by rainfall. The savannah ecosystem of Amboseli is characterized by a five-month-long dry season (June to October), followed by a seven-month period of highly variable rainfall. When baboons were sampled in months with more rainfall, they harbored more within-species genetic diversity and were more likely to share low-abundance strains. We speculate that this elevated strain-level diversity may be a result of interacting with (and eating) more diverse types of vegetation in rainier periods^55^, or by rain-driven activation of dormant microbial populations in the soil^56,57^. The latter explanation may be particularly important for explaining how animals living in different time periods nevertheless can take up nearly identical strains.

In summary, strain sharing patterns in the Amboseli baboon population are not strongly driven by known patterns of social interaction, at least in this temporally and environmentally heterogeneous sample. This result contrasts with the fecal microbiota transplant data, where strain sharing clearly recapitulated transmission pathways. One possible explanation for this difference is that social interactions are not a significant pathway for microbiome transmission in reality, and that host-to-host transmission was detected in the FMT because the transplant method bypassed normal colonization routes. However, other work on this population has reported elevated microbiome similarity among close social partners, both at the species level^2^ and the strain level^36^. In these previous analyses, all samples were collected close in time (i.e., within a two-month period) during a period of relatively little environmental variation. We therefore suspect that a more important explanation relates to the fact that strain-level compositional patterns in microbiomes are the product of both transmission *and* persistence. The distribution of a given strain may reflect its transmission history for a short time after transmission, but in the long term, other ecological processes such as selection, priority effects, and demographic stochasticity will determine whether it persists or goes extinct within each host^58–60^.

Following this reasoning, the best strategy for identifying transmission networks in natural populations may be to focus on recently acquired strains. This approach in turn underscores the importance of longitudinal sampling, which provides the key information needed to identify newly acquired strains. Another reason to sample repeatedly within short time intervals is to minimize the possibility of misclassifying strain sharing events as transmission (e.g., due to independent acquisition of microbes from environmental reservoirs). The genetic similarity threshold recommended for the inStrain pipeline, which we deployed here, is designed to discriminate between strains that diverged as recently as 2.2 years, but this calculation is based on substitution rates in the human gut^26,61^. Bacteria grow and evolve more slowly in many other environments, including soil^62^. In our study, strain sharing across non-overlapping baboon generations was elevated if one or both of individuals in a pair were sampled during a rainy month. If rain revives dormant bacterial populations in environmental reservoirs^56,57^, then these strain sharing events may represent long periods of little evolutionary change between hosts rather than continuous transmission of actively evolving bacterial lineages.

Another conclusion from this study is the value of sampling social networks as completely as possible to obtain information about bacterial distributions at the population level. Our analyses support the idea that some bacterial species are more likely to mirror true transmission networks than others. In the FMT data set, strains shared by true donor-recipient pairs were enriched for bacteria with otherwise limited prevalence in the population, obligately anaerobic bacteria that are unlikely to survive in the environment, and bacteria reported only in human hosts. This result is consistent with reports in several animal populations that socially shared microbial species are enriched for traits associated with host dependence^2,11,12^, as well as theory predicting that reliance on transmission between hosts will select for such traits^9,63^. Our study therefore suggests that filtering out widely shared, generalist strains that may be acquired independently from host-to-host transmission can improve the signal of true transmission networks.

Another consideration is the selection of an appropriate strain profiling method and reference database, especially when working in non-human or non-model systems. We asked whether the main conclusions of our study would hold using an alternative, phylogeny-based strain inference approach. Estimates of strain sharing rates were highly correlated across pipelines, and conclusions based on strain sharing patterns across categories/dyad types were qualitatively consistent. Both pipelines performed well at classifying reads from human-derived metagenomes, but in the baboon data set, StrainPhlAn was limited by its reliance on a reference database in which non-human microbes are still poorly represented. In contrast, inStrain can easily accommodate user-provided genomes, which greatly improved database coverage of the baboon dataset (Figure **S8a**). However, it is still possible that inStrain misses signals of social transmission in the remaining unmapped reads. For this to occur, socially transmitted species would have to be disproportionately baboon-specific (and therefore missing from human-centric databases) and low-abundance (therefore missing from the metagenome-assembled genomes we used to expand the standard database^46^). Whether either explanation is important in this dataset is a question that only additional metagenome- and culture-based assembly of new microbial genomes can help answer.

Finally, our study strongly supports the importance of considering host traits and environmental conditions, beyond social interaction itself. In the Amboseli baboon data set, strain sharing was elevated among individuals who spent their lives in different social groups if they were eating similar diets at the time of sampling. Diet is frequently confounded with sociality in species where social partners forage together, share or steal food, or interact with similar parts of a heterogeneous landscape^19,64,65^. Other characteristics with the potential to affect microbiome composition, such as age and host genetics, are often more similar among social partners as well^20,66^. Studies that simply compare strain sharing within and across social units without considering these additional confounds are at risk of overestimating the contribution of social transmission. More generally, our findings argue that social transmission should not be treated as the default explanation for observations of species *or* strain sharing among interacting individuals.

In summary, strain-resolved metagenomic analyses have clear value for resolving microbiome transmission networks beyond the resolution of species- or genus-level profiling. However, the insights achievable from strain sharing analyses can be improved by careful study design. The proportion of strains shared between individuals is the result of three processes: (i) how many strains they have exchanged via transmission, (ii) how many strains they have independently acquired (e.g. from the environment or other hosts); and (iii) how many of those strains they have both independently retained since transmission occurred. The latter two processes are facilitated by shared environments and can inflate estimates of transmission among social partners. As an alternative approach to coarse estimates of strain sharing rates, we recommend using repeated longitudinal sampling to identify strains that move between hosts within a short period of time. These recently acquired strains are more likely to be reliable indicators of the transmission network, as they have not yet been subjected to extended selection pressures that would alter their distribution in the population. In addition to sampling design, it is also critical to consider ecological, behavioral, genetic, and demographic characteristics that shape microbiome composition, *especially* when those characteristics are more similar among social partners. Importantly, as in the case of our unexpected finding of greater sharing across wetter months–even when temporally separated by years–taking these additional factors into account can also suggest interesting new routes for mapping the transmission landscapes of microbiomes in the wild.

## METHODS

### Fecal microbiota transplant

We re-analyzed publicly available metagenomic data from a study of healthy human donors (n=5) and patients with either recurrent *Clostridium difficile* infection or mild-to-moderately active inflammatory bowel disease (n=8), living in Rome, Italy^38^. Each patient was treated with a fecal transplant from a single donor. Stool samples were collected and sequenced from patients immediately before the transplant and 15-30 days after the transplant. DNA extraction was performed by the original authors using the DNeasy PowerSoil Pro Kit (Qiagen). Libraries were prepared using the Illumina DNA Prep (M) Tagmentation kit and sequenced on the Illumina NovaSeq 6000 platform.

We downloaded raw reads for the fecal transplant study from the European Nucleotide Accession (PRJEB47909) and filtered reads using Trimmomatic^67^, requiring a minimum length of 70 bp and a minimum quality score of 20 within a 4-bp sliding window. Next, we aligned reads to 4,644 species-representative microbial genomes from the Unified Human Gastrointestinal Genome database^68^ using bowtie2^69^. Read counts and mapping statistics are available in Table **S1**. We conducted strain-level population genetic comparisons using the *profile* and *compare* functions of inStrain^26^. Following the recommendation of the developers, we considered a strain to be “present” in a pair of samples if at least 25% of its genome was represented with at least 5x coverage in both samples. We calculated the average nucleotide identity (ANI) of present strains using a microdiversity-aware approach that calls a substitution only when no alleles (major or minor) are shared between the two samples. We considered two samples to share a strain if their strains had 99.999% ANI.

### Sample collection from the baboon field study

The newly generated sequences in this study originated from a population of wild baboons (admixed between *Papio cynocephalus* and *Papio anubis*, with *P. cynocephalus* the majority ancestry) inhabiting the Amboseli basin in southern Kenya. The population has been under continuous study since 1971; the present study includes fecal samples collected between 2007 and 2017. After collection, fecal samples were stored in 95% ethanol at 4°C for up to two weeks, then freeze-dried as follows: 1) All ethanol was evaporated from samples under a fume hood, 2) Tubes were cooled for 30 minutes at -20°C, 3) Tubes were placed in a freeze-dryer (<-50°C, vacuum at 30 millitor). The resulting powders were stored at -80°C.

We selected fecal samples for strain sharing analysis that satisfied one or more of four criteria, focusing our efforts on adult females. First, we selected 20 pairs of fecal samples from female baboons whose lives never overlapped. Second, we selected 23 pairs of fecal samples from baboons living in different social groups at the same approximate time (i.e., collected less than 150 days apart). Note that females do not disperse in this species, meaning that females could not have transferred between distinct social groups prior to sample collection. Third, we selected fecal samples that belonged to close social partners in the same social group, sampled less than four days apart. To identify close social partners, we calculated dyadic sociality indices (DSI, a measure of social bond strength^70^) between females based on all observed grooming interactions between 2007-2017, then sampled 26 unique dyadic pairs from the top quartile of the DSI distribution. Finally, we selected longitudinally collected fecal samples from 22 individuals for whom repeated samples were available within a 120-150 day interval. Note that the same individual could be represented in multiple categories with different partners, but not multiple dyads within the same category (Table **S2)**.

### Metagenome sequencing of baboon samples

Gut metagenomes were generated from the 126 fecal samples selected as described above. First, we extracted microbial DNA using the MoBio PowerSoil Kit with a modified protocol optimized for freeze-dried samples. Specifically, we increased the PowerBead solution to 950 µL/well and incubated the plates at 60°C for 10 minutes after lysis in order to increase the hydration levels of freeze-dried samples and minimize the risk of plate clogging. We prepared libraries using the SeqWell purePlex DNA library prep kit and sequenced the libraries on the NovaSeq X at the University of Chicago DNA Sequencing Facility to a median depth of 34 million read pairs per sample (using a paired end, 150 bp read length design). We removed adapters and filtered raw reads using Trimmomatic^67^, requiring a minimum length of 70 bp and a minimum quality score of 20 within a 4-bp sliding window. Read counts after quality control are available in Table **S3**. Raw data are available on the Sequence Read Archive (accession PRJNA1135081).

As database coverage of non-human metagenomes can be low, we created a custom microbial genome database by supplementing the Unified Human Gastrointestinal Genome (UHGG) database with an additional 2985 genomes assembled directly from the metagenomes of non-human primates, including the baboon population in this study^46^. The full set of genomes was filtered using dRep^71^ to minimize mis-mapping between closely related genomes. Briefly, genomes were clustered into bins based on 95% average nucleotide identity. Representative genomes were selected based on completeness, contamination, and centrality to other genomes in the cluster, with an additional weight using the *–extra_weight_table* flag to favor genomes from non-human primates over those from humans. The final database contained 4712 genomes. The custom database substantially improved read alignment compared to the standard UHGG database (paired t=29.85, p<0.001, Figure **S8a**).

We aligned reads to the custom database using bowtie2^69^ (mapping statistics available in Figure **S8b** and Table **S3**). We conducted strain-level population genetic comparisons using the *profile* and *compare* functions of inStrain. We considered a strain to be present in a pair of samples if at least 25% of its genome was represented with at least 5x coverage in both samples, and we considered samples to be strain sharing if their strains had ≥99.999% ANI.

To compare the major conclusions of the study with an alternative analysis pipeline, we additionally profiled metagenomes at the species level using MetaPhlAn^45^. We extracted clade-specific marker genes using the *extract_markers* function of StrainPhlAn^27^. We used the *strainphlan* function to build species-level phylogenetic trees, requiring each species to be present with 10 or more markers in at least 4 individuals, then extracted the pairwise phylogenetic distances with the *tree_pairwisedists* function of StrainPhlAn. We considered samples to be strain sharing if the normalized phylogenetic distance between them was ≤0.1.

### Behavioral, demographic, and ecological data from the baboon field study

We collected data on the baboons’ diets using point sampling at one-minute intervals within ten-minute focal samples by recording the food type if the individual was feeding during the point sample^72^. We aggregated these data by year and social group, meaning that an individual’s value represents the average diet of her social group in the year of sampling. Comparisons were thus only possible for dyads from different social groups or who never lived at the same time. We generated a Bray-Curtis dissimilarity matrix based on the compositional data using the R package *vegan* and measured the relationship between dietary (dis)similarity and bacterial strain sharing. Daily rainfall values were collected using a rain gauge established at the nearby field camp of the Amboseli Baboon Research Project. Ages of all individuals in our sample were known within a few days as they were followed since birth.

### Taxonomic and functional annotations of shared species

We used GTDB-TK to assign taxonomic classifications to the metagenome-assembled genomes in our custom database based on their sequence similarity to the bacterial and archaeal reference trees available in the Genome Taxonomy Database (GTDB)^73^. We then annotated the oxygen tolerance and known host(s) of each species-representative genomes in the Unified Human Gastrointestinal Genome database using the Genomes Online Database^41^. If all entries in the containing genus were recorded as obligate aerobes, we considered the species aerobic. If all entries in the containing genus were recorded as obligate anaerobes, we considered the species anaerobic. If the genus contained a mixture of obligate aerobes, obligate anaerobes, and/or facultative aerobes or anaerobes, we assigned it the label “mixed”. All genera in the Unified Human Gastrointestinal Genome database had been previously reported in humans at a minimum; some had also been detected in other animals. We thus annotated each entry as “*Homo sapiens* only” or “multiple hosts”.

### Quantification and statistical analysis

We assessed differences in strain sharing between matched and mismatched pairs (FMT dataset) or among categories (baboon dataset) using t-tests and Benjamini-Hochberg correction of p-values. We assessed taxonomic differences between matched and mismatched pairs (FMT dataset) or among categories (baboon dataset) using Fisher’s exact tests, followed by calculation of family-wise log odds ratios if the Fisher’s test was significant. We assessed differences in anaerobic metabolism and host specificity in the FMT dataset using permutation tests, because the filtering criteria imposed on the FMT dataset generated subsets of the full set of strain sharing events and the phenotypes were therefore not independent of the full set or directly comparable to the full set. For the permutation tests, we sampled from the full list of strain sharing without replacement to a size matching the number of strain sharing events detected following a given set of filtering criteria. The p-value represents the proportion of permutations (out of 1000) for which the simulated proportions of anaerobic or host-specific bacteria were as or more extreme as the observed proportions. Information about test statistics, degrees of freedom, and p-values are available in the main text and figure captions.

### Computational Resources

| Software or Database | Reference |
| --- | --- |
| Trimmomatic v.0.39 | Bolger et al. 2014<sup>67</sup> |
| Bowtie2 v.2.5.1 | Langmead and Salberg 2012<sup>69</sup> |
| inStrain v1.3.1 | Olm et al. 2021<sup>26</sup> |
| dRep v3.4.5 | Olm et al. 2017<sup>71</sup> |
| MetaPhlAn v3.0 | Blanco-Míguez et al. 2023<sup>45</sup> |
| StrainPhlAn v4.0.6 | Beghini et al. 2021<sup>27</sup> |
| GTDB-TK v2.3.2 | Chaumeil et al. 2022<sup>73</sup> |
| Unified Human Gastrointestinal Genome database | Almeida et al. 2020<sup>68</sup> |
| ChocoPhlAn vOct22 | Beghini et al. 2021<sup>27</sup> |
| Nonhuman primate metagenome-assembled genomes | Manara et al. 2019<sup>46</sup> |
| Genome Taxonomy Database | Chaumeil et al. 2022<sup>73</sup> |
| Genomes Online Database | Mukherjee et al. 2020<sup>41</sup> |

## Data and code availability

This paper analyzes a combination of existing, publicly available data and newly generated metagenomic sequences. The newly generated sequences will be available on the NCBI Sequence Read Archive upon publication. All original code is available at the Github repository reenadebray/microbiome-strain-sharing. Any additional information required to reanalyze the data reported in this paper is available from the lead contact upon request.

## Author contributions

Conceptualization: RD and JT; Methodology: RD, SEW, CD, EA and JT; Investigation: RD, SEW, CD; Formal Analysis: RD, CD; Resources: EA, JT; Data Curation: RD, SEW, CD, EA, JT; Writing – Original Draft: RD, JT; Writing – Review & Editing: SEW, CD, EA, JT; Visualization: RD; Supervision: EA, JT; Project Administration: EA, JT; Funding Acquisition: EA, JT.

## Acknowledgments

We gratefully acknowledge the support of the National Science Foundation and the National Institutes of Health for the majority of the data represented here, currently through R01AG071684, R01AG075914, and R61AG078470, as well as prior funding from the National Science Foundation that supported microbiome data generation in Amboseli, including IOS-1053461 and DEB-01840223. Current support for field-based data collection also comes from the Max Planck Institute for Evolutionary Anthropology, and we thank Duke University, Princeton University, and the University of Notre Dame for financial and logistical support. In Kenya, our research was approved by the Wildlife Research Training Institute (WRTI), Kenya Wildlife Service (KWS), the National Commission for Science, Technology, and Innovation (NACOSTI), and the National Environment Management Authority (NEMA). We also thank the University of Nairobi, the Kenya Institute of Primate Research (KIPRE), the National Museums of Kenya, the members of the Amboseli-Longido pastoralist communities, the Enduimet Wildlife Management Area, Ker & Downey Safaris, Air Kenya, and Safarilink for their cooperation and assistance in the field. Particular thanks go to the Amboseli Baboon Project long-term field team (R.S. Mututua, S. Sayialel, J.K. Warutere, I.L. Siodi, I.L., and L. Musembei), and to T. Wango and V. Oudu for their assistance in Nairobi. The baboon project database, Babase, is expertly managed by N. Learn, J. Gordon, and W. Wilbur. Database design and programming are provided by K. Pinc. This research was approved by the IACUC at Duke University and the University of Notre Dame, and the Ethics Council of the Max Planck Society and adhered to all the laws and guidelines of Kenya. For a complete set of acknowledgments of funding sources, logistical assistance, and data collection and management, please visit http://amboselibaboons.nd.edu/acknowledgements/.

