## Supplementary Figures for "Shared environments complicate the use of strain-resolved metagenomics to infer microbiome transmission"

Supplementary material for ‘Shared environments complicate the use of strain-resolved metagenomics to infer microbiome transmission’

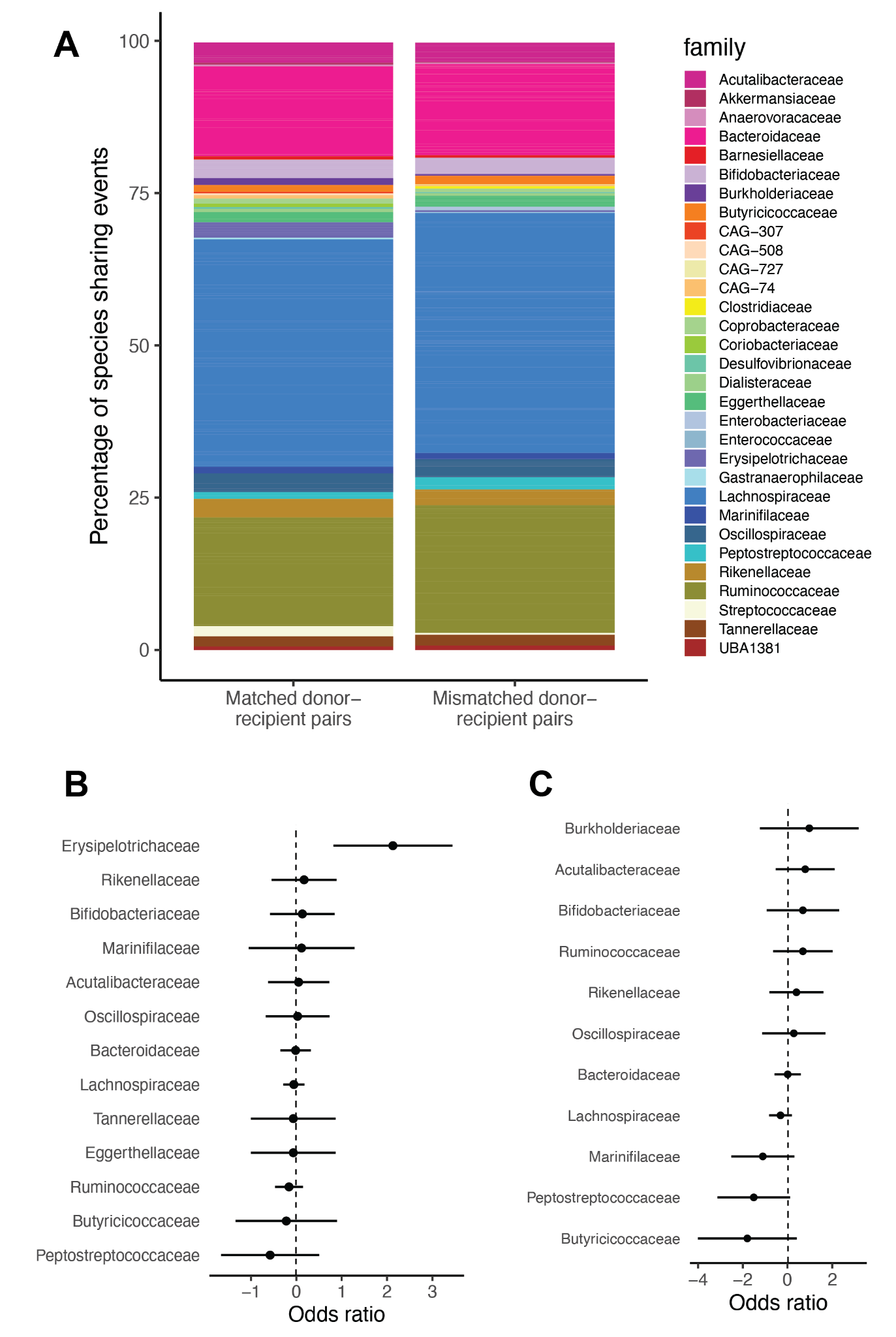

**Figure S1. Bacterial taxa shared between donors and recipients of fecal microbiota transplants. (a) Family-level structure of bacterial taxa that were shared at the species level between subjects.** Bar chart of species (aggregated into families) that exhibit species sharing between matched (left) or mismatched (right) pairs. Colors reflect different bacterial families and are scaled to represent the relative proportions of strain-sharing events in each category**. (b) Species-level sharing of families among matched pairs relative to mismatched pairs.** The log_2_ odds ratio represents the relative likelihood that bacterial families were shared at the species level between matched donor-recipient pairs, compared to their species sharing rates between mismatched pairs. **(c) Strain-level sharing of families among matched pairs relative to mismatched pairs.** The log_2_ odds ratio represents the relative likelihood that bacterial families were shared at the strain level between matched donor-recipient pairs, compared to their strain sharing rates between mismatched pairs. Error bars represent 95% confidence intervals. Error bars overlapping 0 (dashed line) are not significantly enriched or depleted in matched pairs.

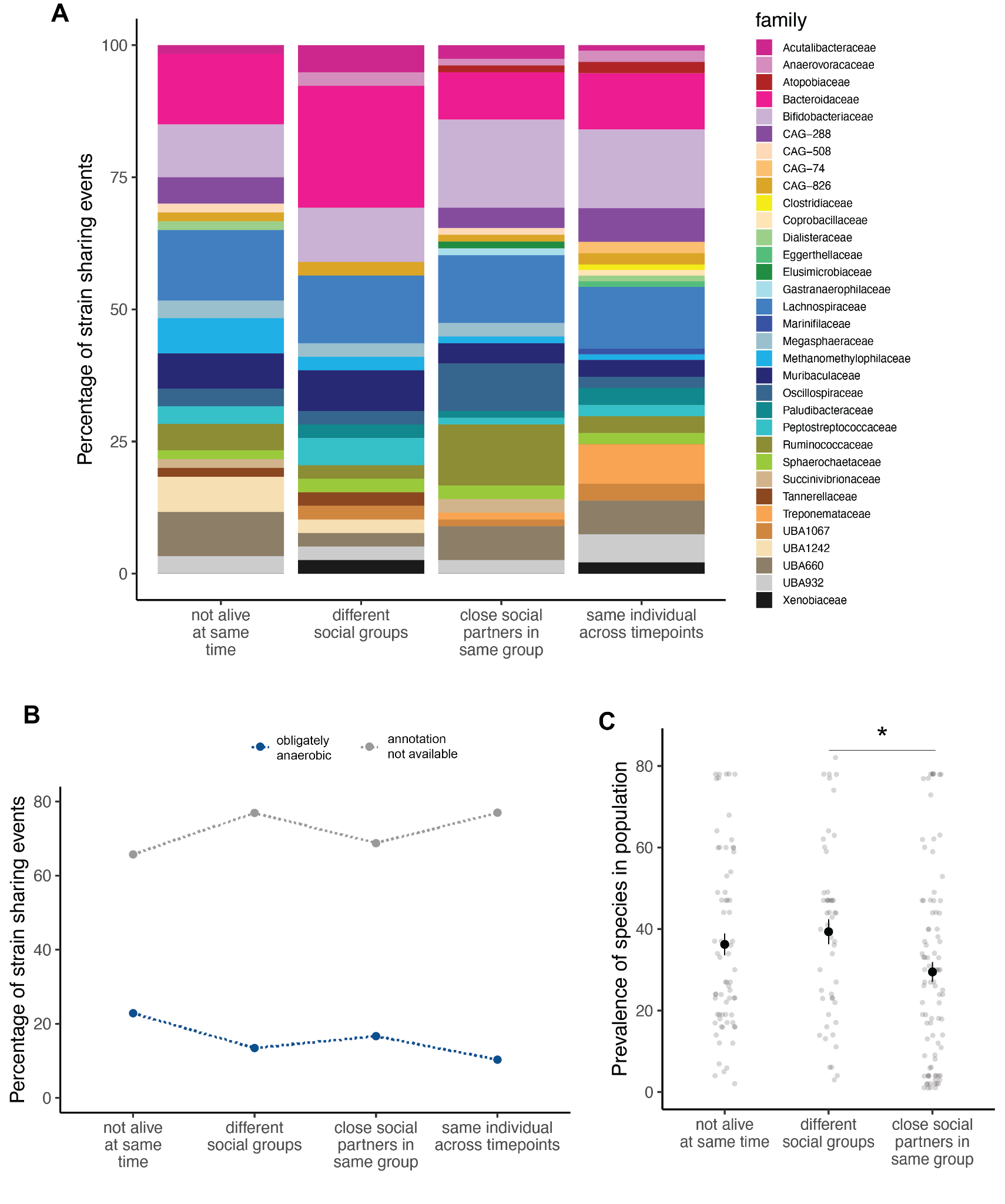

**Figure S2. Taxonomic and functional characteristics of bacterial taxa in which strain sharing was detected between baboons in a wild population. (a) Family-level structure of bacterial taxa that were shared at the strain level between individuals.** Bar chart of species (aggregated into families) that exhibit strain-sharing between pairs of baboons, based on a 99.999% ANI threshold. Colors reflect different bacterial families and are scaled to represent the relative proportions of strain-sharing events in each category. **(b) Anaerobic metabolism across varying definitions of transmission.** Strain sharing events were annotated based on phenotype information available in the Genomes OnLine database. **(c) Population prevalence of bacterial species.** Each strain sharing event was annotated according to the prevalence of the species to which it belonged, excluding later longitudinal samples to avoid double-counting individuals. The y-axis represents the number of baboons in which the species was detected, out of a total n=93.

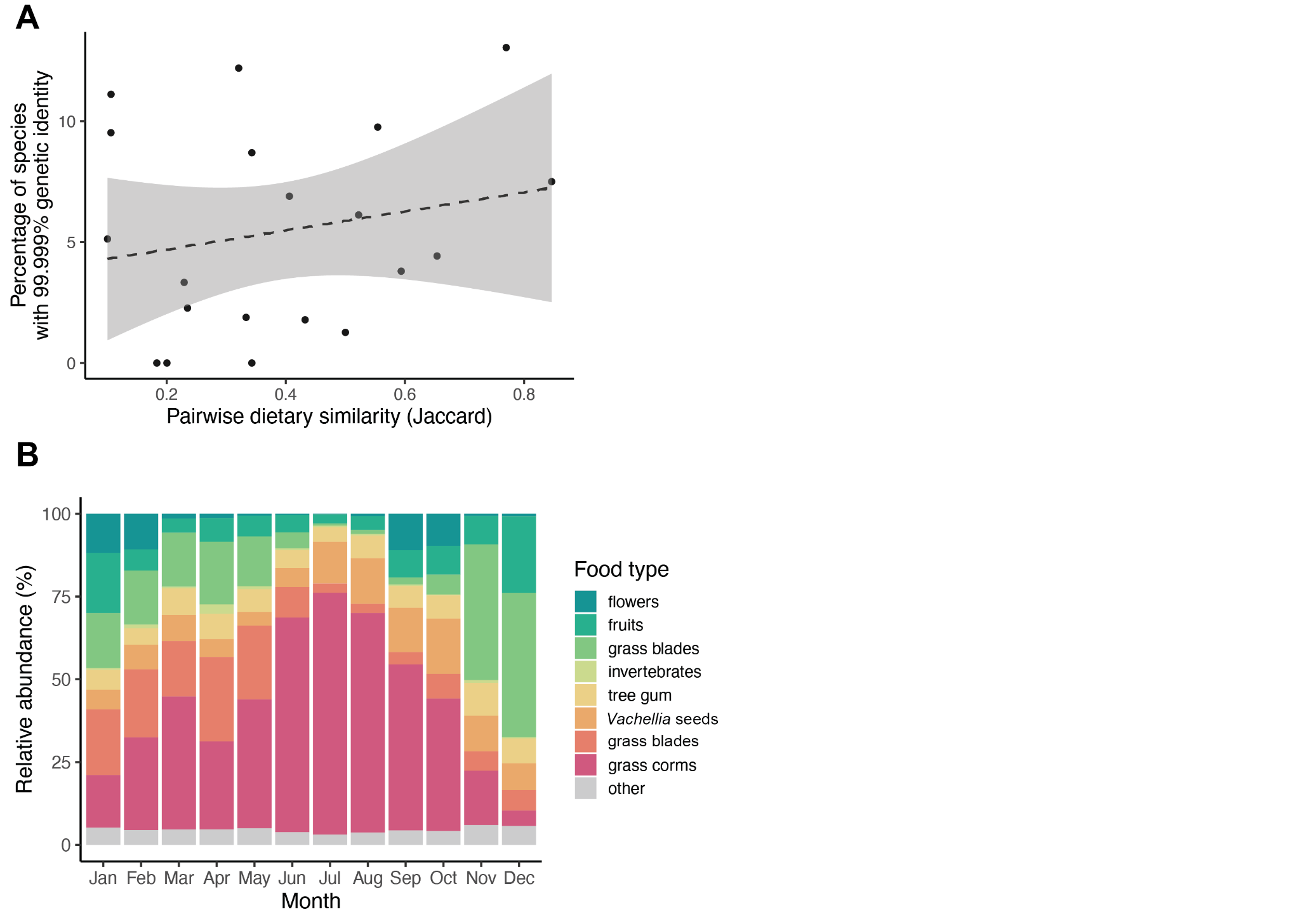

**Figure S3. Diet and seasonality in Amboseli. (a) Dietary similarity between social groups.** Dietary similarity was calculated using the Jaccard similarity index based on the dietary compositions of each pair of baboons that lived in different social groups at similar times, aggregated by month. **(b) Variation in diets throughout the year.** Colors reflect different foods consumed by the baboons between 2007-2017 and are scaled to represent the relative proportions consumed in each month.

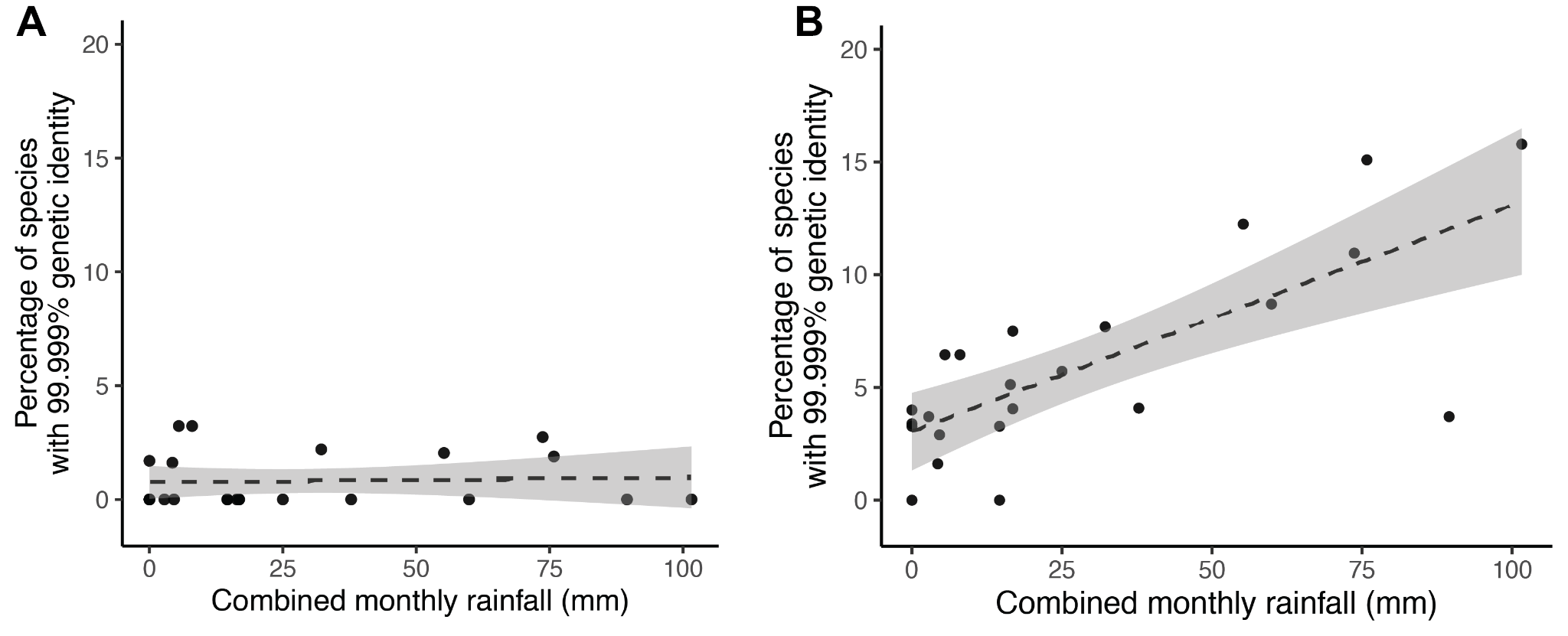

**Figure S4. A consensus-based approach does not detect an effect of rainfall on strain sharing.** Rainfall was measured daily using a rain gauge and summed by month for samples taken from baboons that lived at different times. The y-axis represents strain sharing based on the consensus average nucleotide identity (conANI) value calculated by inStrain. This metric considers two genomes to differ at a given site if their consensus alleles are different (i.e., it ignores minor allele sharing). **(b) Microdiversity-aware approach (reproduced from Figure 4b).** This metric considers two genomes to differ at a given site if they do not share any alleles at that site (major or minor).

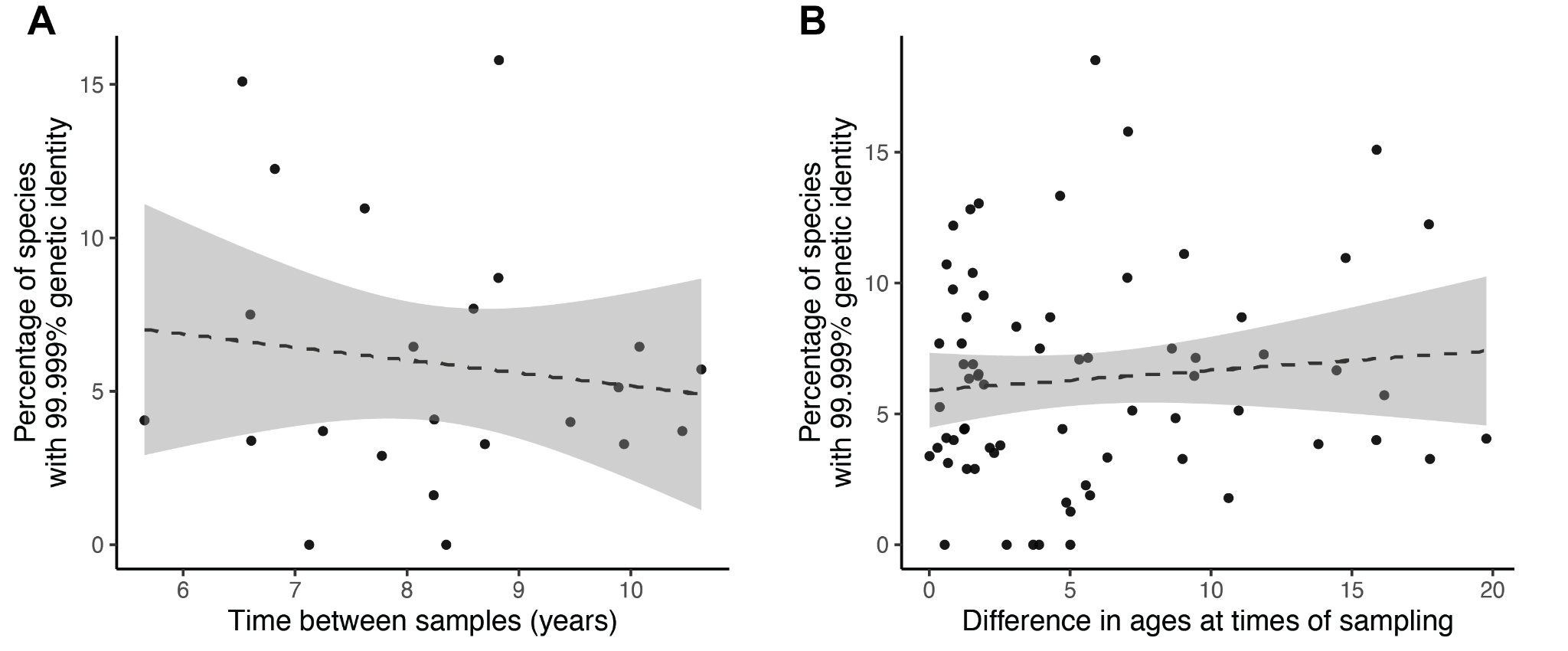

**Figure S5. Host characteristics that did not predict strain sharing rates. (a) Number of years between samples.** Plot includes only baboon pairs that lived at different times, as the remaining dyad types were intentionally sampled within short time periods. (**b) Difference in ages at times of sampling.** Chronological ages were known with high confidence because all subjects were born in regularly censused study groups.

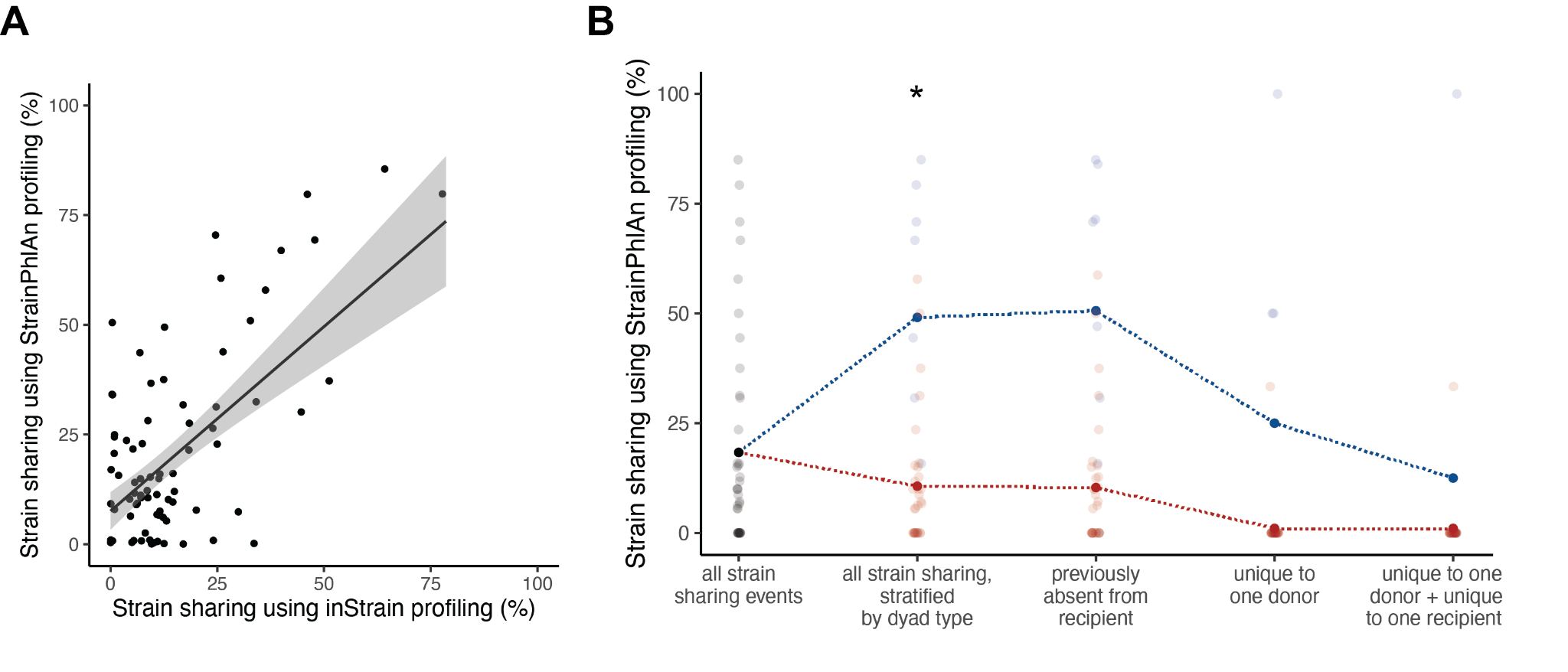

**Figure S6. Strain sharing analysis of FMT dataset with StrainPhlAn pipeline. (a) Correlation between inStrain and StrainPhlAn estimates of strain sharing**. Each point represents a donor-recipient pair (either matched or mismatched) in the FMT dataset. Samples with fewer than three shared species (i.e., the denominator in the calculation of strain sharing rates) are excluded from this visualization. **(b) Strain sharing across varying definitions of transmission.** The percentage of strain sharing events among matched donor-recipient dyads (blue) and all other comparisons (red) is shown following serially more stringent filtering criteria. Asterisks represent significant differences between matched and mismatched cohorts based on t-test and Benjamini-Hochberg correction: (***) p<0.001; (**) 0.001≤p<0.01; (*) 0.01≤p<0.05.

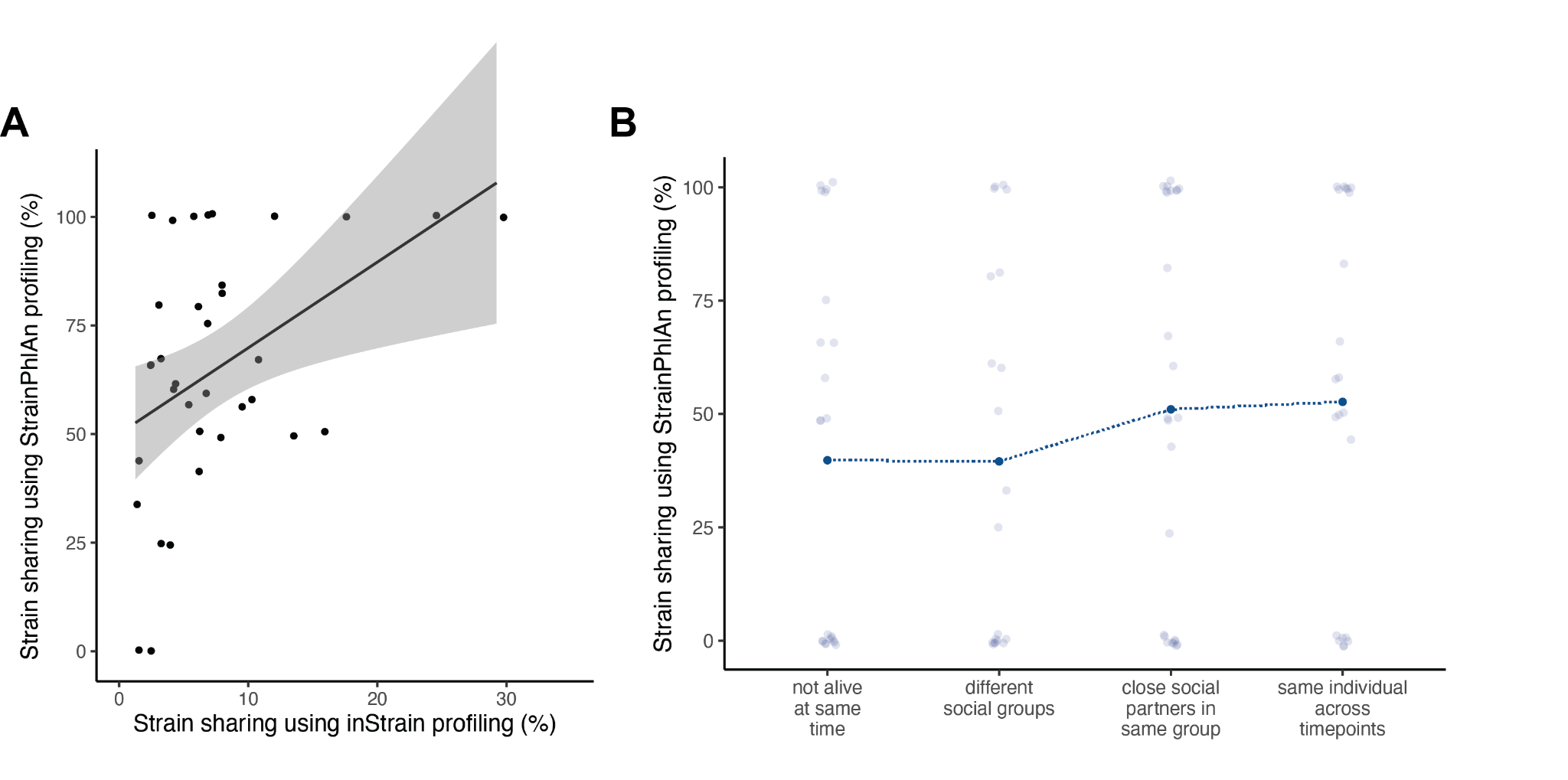

**Figure S7. Strain sharing analysis of baboon dataset with StrainPhlAn pipeline. (a) Correlation between inStrain and StrainPhlAn estimates of strain sharing**. Each point represents a donor-recipient pair in the FMT dataset. Samples with fewer than three shared species (i.e., the denominator in the calculation of strain sharing rates) are excluded from this visualization. **(b) Strain sharing rates across dyad types.** The percentage of shared strains between each dyad (≤0.1 normalized phylogenetic distance).

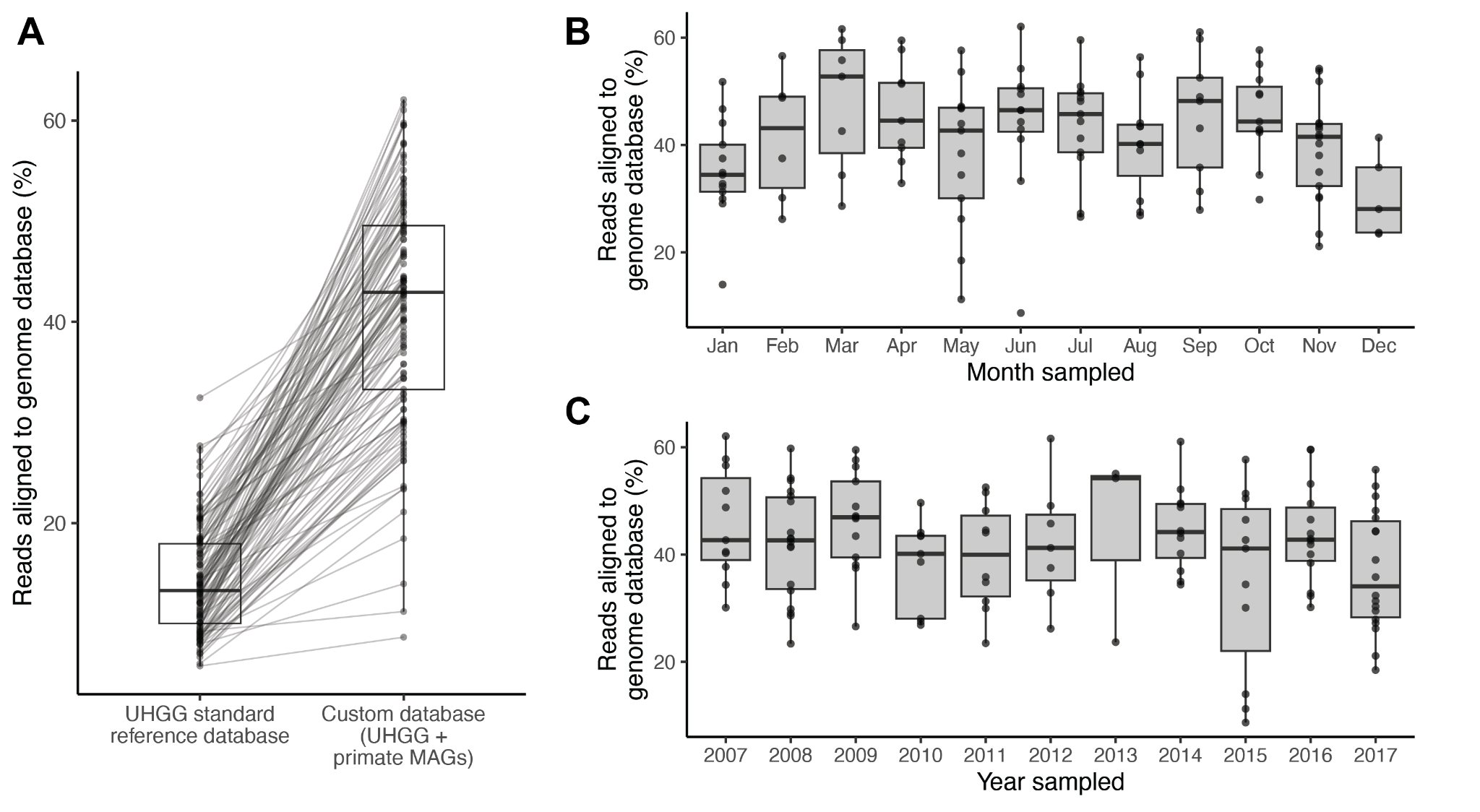

**Figure S8. Database coverage in baboon gut metagenome dataset. (a) Database customization with metagenome-assembled genomes.** Percentage of reads after quality filtering that aligned to microbial genomes in either the Unified Human Gastrointestinal Genome (UHGG) database or the UHGG database plus metagenome-assembled genomes from non-human primate fecal samples. **(b) Database coverage by month that the sample was collected.** Database coverage varied slightly by sampling month (ANOVA, p=0.053), with a tendency for samples from the dry season to map better than samples from the wet season. **(c) Database coverage by year that the sample was collected**. Database coverage did not vary by year.
